# Green turtle satellite telemetry: identifying spatial movements, residency patterns, and habitat associations in Aotearoa New Zealand

**DOI:** 10.64898/2026.10.07.757418

**Authors:** Karen L. Middlemiss, Michelle Boyle, Giorgia Vattiato, James Chatterton, Adam Naylor, Celine Campana, Andrew Christie, Isabella Jansen

**Author notes:** **CORRESPONDING AUTHOR:** Karen L. Middlemiss.

## Abstract

Sea turtle conservation relies largely on the identification of ecologically significant marine areas. Green sea turtles (*Chelonia mydas*) are resident in Aotearoa New Zealand waters, yet much of their spatial ecology remains undefined. To address this, satellite telemetry was used to map juvenile and sub-adult green sea turtle spatial movements, residency patterns, and habitat associations. Our findings provide the first evidence of individual fine-scale habitat use and long-term residency in New Zealand waters. Data will contribute to regional and global efforts to identify ecologically significant sea turtle habitat, including Important Marine Turtle Areas (IMTAs) and Blue Corridors for Turtles (BC4T), and help advance conservation efforts.

## 1. INTRODUCTION

Coastal harbours are globally recognised as important post-settlement foraging habitats for juvenile and sub-adult green sea turtles (Arthur et al. 2008). In northern New Zealand, several harbours contain suitable habitat characteristics for this species (Morrison et al. 2009). Rangaunu Harbour, for example, is a 115 km^2^ shallow-water system supporting seagrass meadows (*Zostera muelleri*), mangrove forests (*Avicennia marina*), and diverse fish and invertebrate communities (Kerr 2009; Morrisey et al. 2007; Morrison et al. 2014; Morrison et al. 2009). Ecologically, it is comparable to nearby Pārengarenga Harbour, and many similar seagrass habitats occur elsewhere in the North Island of New Zealand, including Manukau Harbour (Morrison et al. 2009).

Green sea turtles typically inhabit tropical to sub-tropical waters in temperatures above 15° C (Read et al., 1996). However, in a 2023 a green sea turtle in Australian waters spent a month in mean Sea Surface Temperatures (SST) of 12° C and as low as 8.5° C highlighting individual variation in thermal tolerances (K. Middlemiss, pers. comm.). Sea temperatures in northern New Zealand typically sit above 15° C year-round (Godoy, 2008). Seasonal temperature variation influences sea turtle movement and depth use, potentially through behavioural thermoregulation (Crear et al., 2016).

Despite the availability of apparently suitable habitat, and the regular occurrence of green sea turtles in New Zealand waters, there is no published fine-scale data quantifying and mapping habitat types to spatial movements or residency patterns for individual turtles. Previous distribution studies have relied largely on stranding and sighting records (Godoy et al. 2016). These knowledge gaps constrain our understanding of green turtle spatial ecology. Data on persistent habitat use and site fidelity are fundamental to conservation planning, spatial management, and mapping of ecologically significant habitats across migratory ranges. This includes global initiatives such as Important Marine Turtle Areas (IMTAs) and Blue Corridors for Turtles (BC4T). Here, we present findings from the first published satellite telemetry study of green sea turtles in New Zealand characterising habitat associations, residency patterns, and seasonal spatial movement patterns for individual turtles in relation to temperature gradients. Our aim is to report important initial findings from our research to begin informing protection of green sea turtle habitat in New Zealand and to support international conservation efforts.

## 2. MATERIALS AND METHODS

Eleven rehabilitated green sea turtles were satellite-tagged and released in Rangaunu Harbour (34.9482° S, 173.2702° E, Northland, New Zealand) between March 2025 - March 2026, near their original stranding locations. Except for one turtle that was released in Taupō Bay where it stranded. It is not known if they had strong associations with the area prior to stranding. All turtles were rehabilitated at Auckland Zoo and Sealife Kelly Tarlton’s. Size classes were juvenile (n=6) and sub-adult (n=5) with curved carapace lengths (CCL) ranging from 47-80 cm and weights between 13-62 kg^-1^. Juvenile size was accepted at 30-59 cm CCL and sub-adults 60-90cm CCL based on Limpus and Chaloupka (1997).

### Satellite tagging

Turtles were fitted with F385 (sub-adult) or SPLASH10-F427 (juvenile) Fastloc® GPS tags (Wildlife Computers Incorporated) using attachment methods described in Hays and Hawkes (2018). Standard tag transmission times, based on estimated battery life, were achieved at were 6- and 12-months respectively for F427 and F435. Tags were configured for green shell turtles using pre-programmed configurations by Wildlife Computers. Fastloc® ARGOS data was analysed using data processing from a minimum of 5 satellite passes to ensure a high level of location accuracy estimated to be within 10-20 m. GPS snapshot information was collected from the tags including SST, depth and location coordinates.

### Habitat mapping in Rangaunu Harbour

Turtle habitat associations within Rangaunu Harbour were evaluated quantitatively using seven mapped habitat classes: channel, mangrove, saltmarsh, channel margin, bare sediment, sparse Seagrass, and dense seagrass. Other harbours were not analysed for habitat types due to resource constraints. Habitat classes represented the broad benthic and intertidal environments used in subsequent telemetry analyses, whereas depth was analysed separately as a continuous environmental variable. The habitat map was developed from satellite imagery, existing authoritative habitat datasets, integrating Sentinel-2 multispectral imagery, LiDAR data, NDVI analyses, object-based image segmentation, supervised classification, aerial imagery interpretation, and existing expert review (Kerr 2009). Mangrove and saltmarsh habitats were derived from the Northland Regional Council (NRC) Intertidal Vegetation Mapping Programme (Kerr 2015) and subsequently reviewed against contemporary imagery to verify their spatial extent. Subtidal habitat classes (seagrass, bare sediment, and channels), were delineated specifically for this study through remote sensing and image interpretation techniques.

### Data Analysis

High-use areas were identified using kernel density estimation (KDE) of 12,962 georeferenced turtle detections from 11 tracked green sea turtles. Individual KDE surfaces were standardised to give each turtle equal contribution and averaged to produce a population-level utilisation surface. Core-use areas were defined as the highest 2% (98^th^ percentile contour) of values in the population-level KDE surface. Additional harbours of interest were identified from a heatmap of all detections pooled across turtles, without standardisation or averaging, to capture harbours of interest not captured by the population-level KDE (Figure 1).

**Figure 1.**
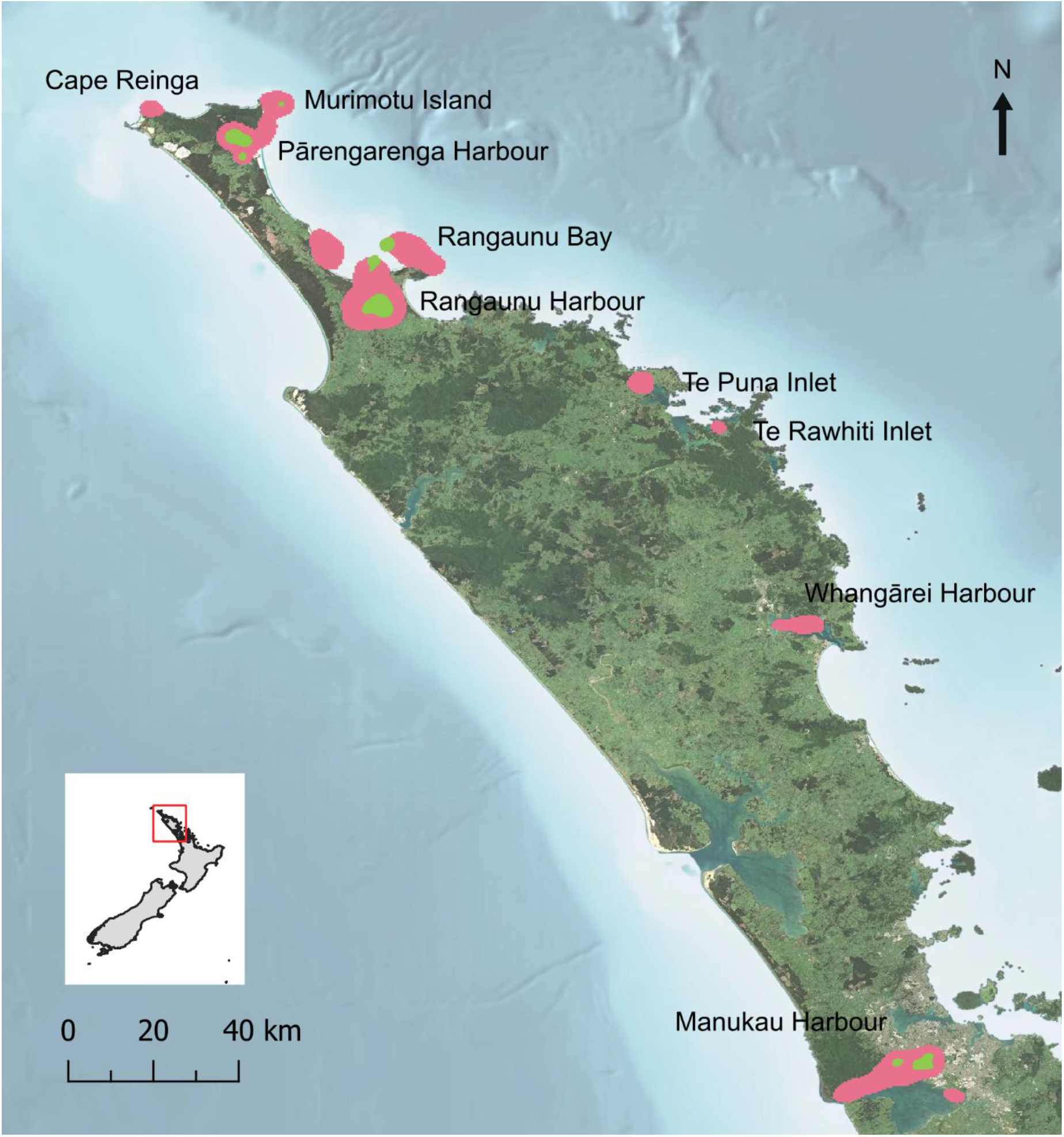
High-use turtle areas across the study region. Green polygons represent areas where turtles spent 30-100 days. Pink polygons indicate additional areas of interest where turtles spent < 30 days.

Within Rangaunu Harbour, habitat use was quantified as the proportion of detections in each habitat class. Depth and temperature were analysed separately, with daily satellite SST variability calculated as the absolute difference between consecutive observations and maximum depth summarised by month and season. Austral seasons were defined as: spring (September – November), summer (December – February), autumn (March – May) and winter (June – August).

Habitat persistence was assessed using consecutive detections within the same habitat and with detections separated by ≤12 h treated as a single residence bout. The 12 h threshold was based on the empirical distribution of detection intervals (median = 2.64 h; 90% ≤6.74 h; 95% ≤10.02 h). This retained 96.3% of consecutive intervals while reducing the likelihood that unobserved habitat transitions occurred between detections. Residence duration was calculated as the elapsed time between the first and last detection within each bout; bouts containing fewer than two detections were excluded.

The probability that the subsequent detection occurred in the same habitat was modelled using a binomial GLMM, with habitat as a fixed-effect and turtle identity as a random intercept. Residence duration was analysed separately using a Gamma GLMM and the same fixed and random effects. Estimated marginal means from the respective models provided habitat-specific probabilities of remaining in the same habitat and mean residence times, respectively, with 95% confidence intervals. Seasonal differences in residence duration were summarised using median residence interval per season.

## 3. RESULTS

### Spatial distribution and habitat use

All turtles remained within North Island coastal waters, with three persistent high-use areas identified in Rangaunu Harbour and Bay, Pārengarenga Harbour, and Manukau Harbour (Fig. 1). Of turtles detected in these areas, 90% remained for a continuous period of >100-days. Other areas visited, but less closely associated with, included Murimotu Island, Te Puna Inlet, Te Rawhiti Inlet, Whangārei Harbour and Cape Reinga (Fig.1).

Spatial use varied among individual turtles, with the strongest site fidelity shown in Rangaunu Harbour (52.5%), followed by Rangaunu Bay (13.6%), and Pārengarenga Harbour (1.2%), while other individuals ranged more widely exploring vast distances of coastline (Fig. 2). This included one turtle that swam over 3000 km exploring the west coast coastline as far south as Wellington before returning to Rangaunu Harbour (Fig. 2A). Another travelled over 1200 km from Taupō Bay south along the east coast to Thames, in the Hauraki Gulf region, before returning north, spending time in Whangārei Harbour and subsequently settling in Rangaunu Harbour (Fig 2D). The median daily movement patterns of sub-adult turtles, calculated as the total straight-line distance between successive locations divided by the total number of tracked days, was nearly two-fold of the juveniles at 19.4 (range 7.3–75.5) km/day and 10.0 (range 1.5–51.9) km/day, respectively.

**Figure 2.**
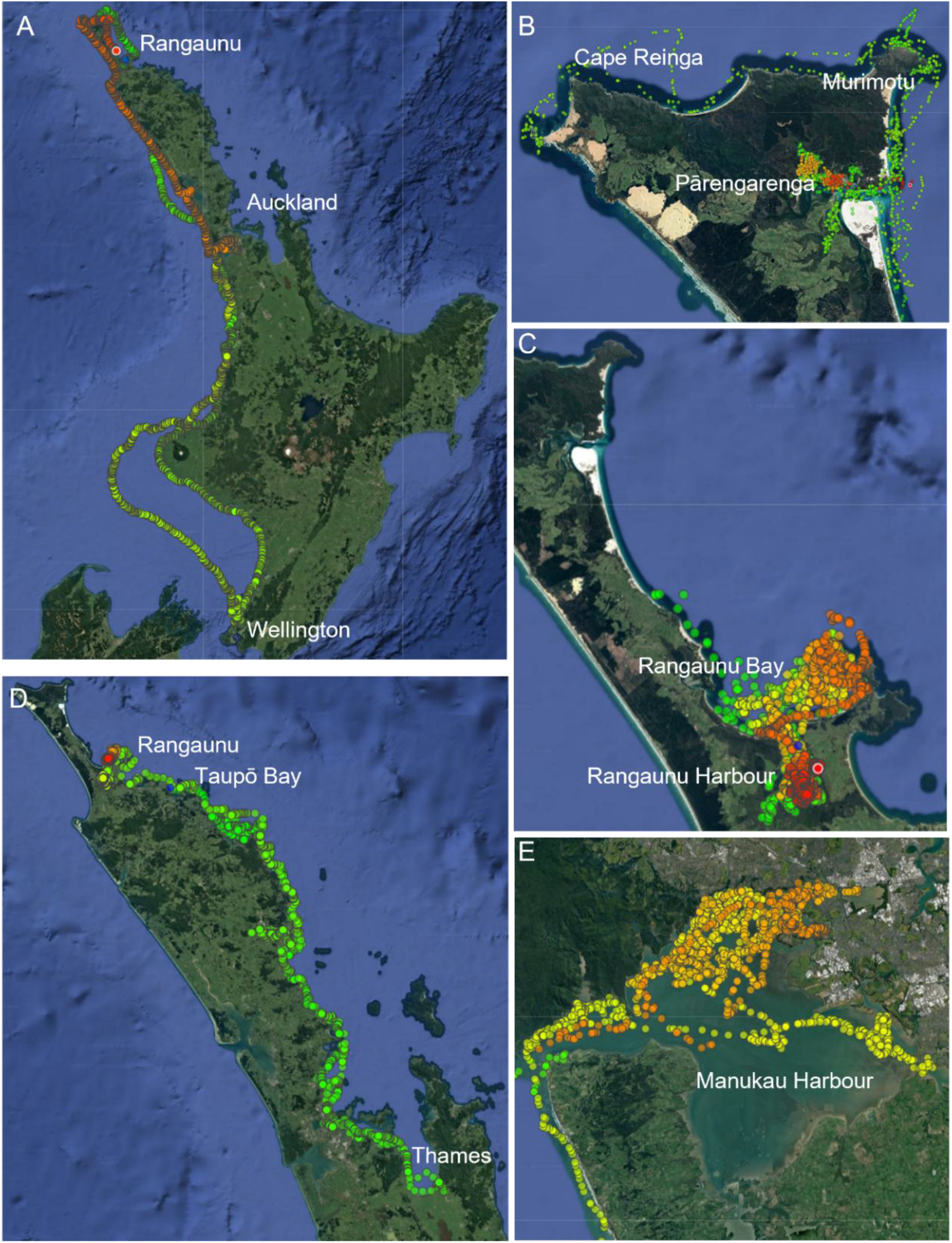
Movement patterns represented by five seperate green sea turtles showing variation in habitat use and movement patterns in North Island, New Zealand coastal areas. (A) Rangaunu to Wellington (B) Pārengarenga Harbour and Murimotu Island, (C) Rangaunu Bay and Harbour, (D) Taupō Bay to Thames, and (E) Manukau Harbour. Colours represent a chronological sliding scale of GPS coordinates from green (first) to red (most recent).

Within the Rangaunu Harbour and Rangaunu Bay hotspots, kernel density analyses identified three distinct high-use zones (Fig. 3A). Seagrass and channel habitats dominated these areas, and habitat use varied between turtles (Fig. 3B). Notably, turtles were minimally associated with seagrass habitat on the western side of Rangaunu Harbour, with detections equivalent to 6.6% versus 93.4% of all seagrass records on the eastern side (Fig. 3B).

**Figure 3.**
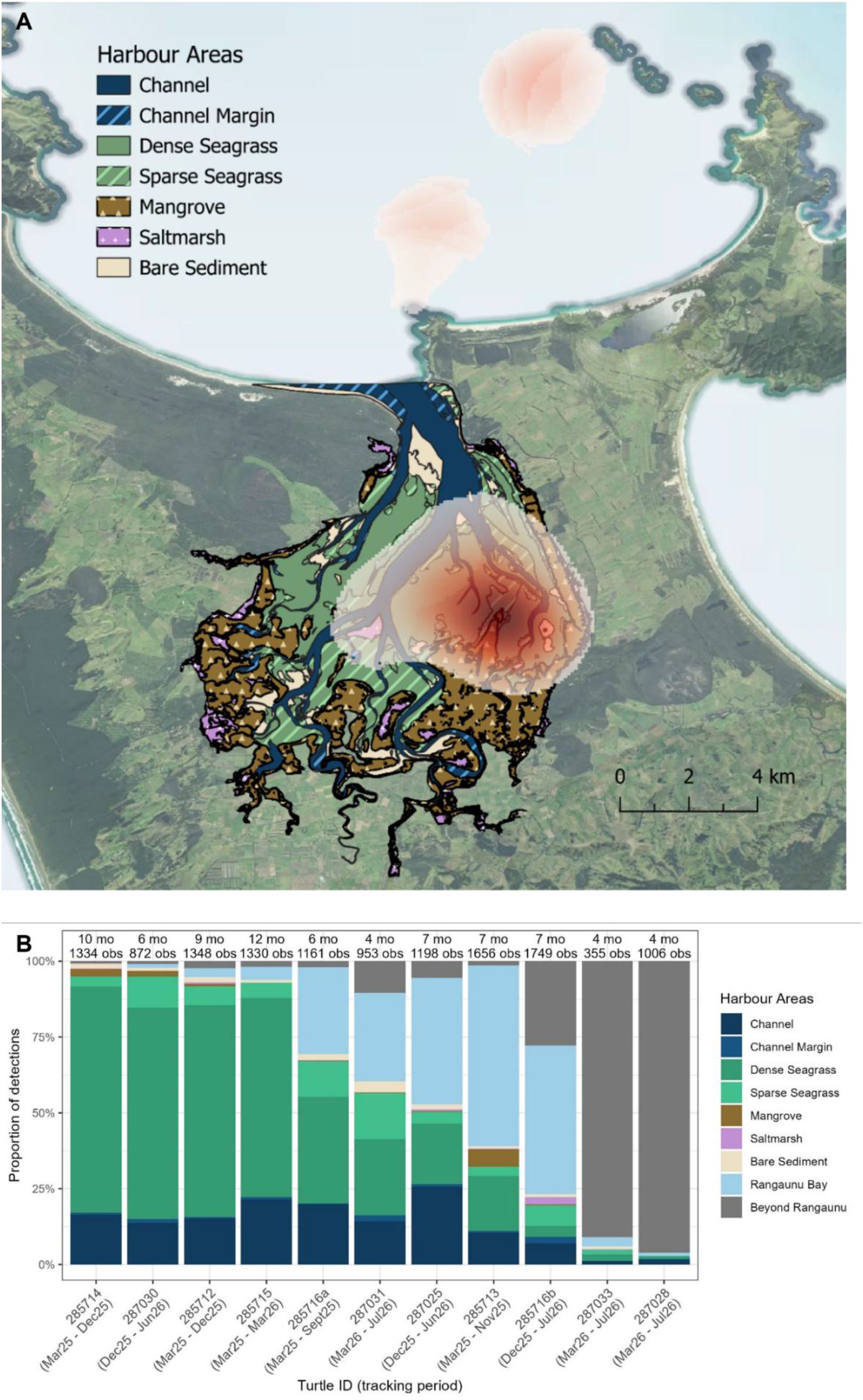
(A) Spatial distribution of Rangaunu Harbour and Rangaunu Bay habitat classes (coloured and patterned polygons), and relative detection density (overlaid kernel density estimate (KDE) heatmap). (B) Proportional distribution of detections in Rangaunu Harbour habitat types, with x-axis labels indicating the turtle ID and tracking period. Numbers above each bar indicate the tracking duration (months) and total number of detections for each individual.

### Habitat fidelity and residence patterns

Habitat fidelity differed among habitat types, with turtles showing the strongest association to dense seagrass (estimated probability = 0.760, 95% CI: 0.741–0.778; n = 4,425 detection pairs). Fidelity was lower in channel habitats (0.438, 95% CI: 0.410–0.468; n = 1,737) and sparse seagrass (0.368, 95% CI: 0.331–0.407; n = 763).

Residency analysis showed a similar pattern, with turtles remaining in dense seagrass for longer than other habitats. The Gamma mixed-effects model estimated mean residence times of 14.2 h (95% CI: 12.5–16.0 h; n = 784 residence bouts) in dense seagrass, compared with 6.1 h (95% CI: 5.3–7.0 h; n = 383 residence bouts) in channels, representing an approximately two-fold difference in residence time. Sparse seagrass had a similar estimated residence time to channels (5.8 h; 95% CI: 4.8–6.9 h; n = 153 residence bouts), with overlapping confidence intervals and little evidence of a difference from channels (*p* = 0.58). Estimates for mangrove, saltmarsh, channel margin and bare sediment habitats were less certain because of limited number of residence bouts (10–25 observations in each) and should therefore be interpreted cautiously.

### Seasonal variation in depth behaviour and SST

SST variability differed among locations, and within the Rangaunu area where SST was lowest in Rangaunu Bay (median absolute daily change = 0.20° C), compared to habitats within Rangaunu Harbour which showed greater variability (median absolute daily change = 0.60° C). Depth and SST data are provided as supplementary material in Fig. S1. Locations outside the Rangaunu area experienced the largest SST changes. The maximum observed daily change was 4.95° C, from approximately 16° C to 11° C, recorded in Manukau Harbour. This temperature shift coincided with a turtle leaving Manukau harbour and travelling directly back to Rangaunu Harbour over a 23-day period, where the SST was around 16° C. Substantial SST variation also occurred while turtles remained within the same broad habitat types, particularly dense seagrass (maximum daily change = 3.13° C) and channel habitats (maximum daily change = 3.83° C).

Depth data showed clear seasonal variation in maximum depth. Greatest depths were recorded during autumn, when turtles had the greatest median maximum depth (10 m) and widest distribution of depths. In contrast, spring was associated with the shallowest depth use (median maximum depth = 3.5 m). Summer and winter showed intermediate depth distributions, although winter exhibited greater variation than summer. Seasonal differences in depth use were not accompanied by major shifts in habitat associations. Turtles remained primarily associated with dense seagrass throughout the year (∼56–58% of observations), with channels the second most frequently used habitat type. Depth use varied within these habitats. In summer months, observations in dense seagrass occurred at relatively shallow depths (median maximum depth = 3.5 m), whereas channel habitats were associated with deeper depths (median maximum depth = 13.5 m). Channel use slightly increased during winter, but turtles did not shift into other habitat types, suggesting that seasonal changes in diving behaviour occurred within the existing habitat mosaic. Habitat occupancy duration also showed limited seasonal variation. Median residence intervals were 5.2 h in summer and 6.2 h in winter, indicating slightly longer periods of continuous habitat use during winter but no major seasonal change in short-term habitat persistence.

## 4. DISCUSSION

Our study provides the first insight into residency and fine-scale habitat use of individual green sea turtles in New Zealand. Data showed that turtles exhibit persistent and predictable use of discrete coastal habitats, connectivity among coastal areas, and strong associations with dense seagrass meadows and harbour channels in Rangaunu Harbour. This is consistent with the species’ benthic feeding ecology (Arthur et al. 2008), and provides evidence that New Zealand’s temperate seagrass ecosystems support sustained habitat use by green sea turtles. The frequent use of harbour channels, perhaps as navigation corridors or resting areas along bathymetric features (Limpus & Coffee 2019), together with seasonal variation in depth-utilisation patterns, identified predictable high-use of spatially discrete zones within Rangaunu Harbour. This generated predictable high-use areas that could contribute to regional turtle conservation strategies as well as current global initiatives to identify ecologically significant turtle habitats, including Important Marine Turtle Areas (IMTAs) and Blue Corridors for Turtles (BC4T).

Individual variation in turtle movement and residency patterns suggest two contrasting habitat-use strategies: long-term occupancy of localised coastal habitats and extensive movements connecting distant regions. Around two thirds of turtles showed strong site fidelity to discrete zones within the Rangaunu Harbour and Rangaunu Bay, while others undertook long-distance coastal movements, indicating connectivity between foraging areas, but often returning to Rangaunu. This highlights the importance of the area, even for turtles not previously known to have close associations. Notably, the turtle which completed a return journey of over 3000 km from Rangaunu Harbour to Wellington, only visited one additional harbour, Manukau, perhaps suggesting a degree of familiarity with, or preference for, specific foraging habitat. We recommend future research to characterise seagrass species found in all harbours visited by green turtles, and that are known to be favoured by green sea turtles (Read & Limpus 2002). Expanding habitat mapping and quantifying habitat availability across New Zealand harbours would improve understanding of green turtle habitat selection and support conservation measures, including the protection and restoration of vital seagrass meadows.

While juvenile and sub-adult turtles showed similar exploratory behaviour outside of the Rangaunu area, median daily movement patterns of sub-adult turtles were double that of juveniles. These differences between size classes are likely related to ontogenetic shifts in dietary preference and feeding strategies between size classes as evidenced in other research on green sea turtles (Pillans et al. 2022). Seasonal changes in depth use without major shifts in habitat associations, suggest that turtles adjust aspects of their behaviour in response to seasonal environmental variation while remaining within established habitat networks. This pattern may reflect behavioural thermoregulation, consistent with findings reported in Crear et al. (2016).

Our results identify Rangaunu, Pārengarenga, and Manukau Harbours as potential IMTAs and highlight key developmental foraging habitats for green sea turtles, and connections between those areas. In Rangaunu Harbour, turtles primarily spent their time in seagrass habitat, and channels; the latter perhaps as navigation corridors or resting areas along bathymetric features. Turtles were almost entirely associated with habitat on the east side of Rangaunu Harbour, and this combined with seasonal shifts in depth behaviour creates predictability in turtle presence/absence in these areas that can inform targeted conservation measures. Future research should investigate the ecological drivers of habitat selection in the eastern side of Rangaunu Harbour, including seagrass distribution, sediment characteristics, and channel hydrodynamics. Expanding our satellite tracking programme to confirm these patterns across a larger sample of turtles will also strengthen findings. Our initial findings have highlighted the conservation importance of New Zealand harbours to migratory green sea turtles, particularly those with seagrass meadows, and provided evidence of the need to incorporate these areas into regional and international spatial planning and conservation strategies.

## 5. ETHICS AND INTEGRITY STATEMENTS

### a. Data Availability

The authors intend to make the data publicly available through Blue Corridors for Turtles (BC4T).

### b. Ethics and Permit Approvals

All research was permitted under the Wildlife Act (1953) with approval from the Department of Conservation Animal Ethics Committee (AEC-482).

### c. Use of Artificial Intelligence

The authors drafted original text and at the final stage minimally used Microsoft Co-Pilot to assist with refining language and grammar and improve audience readability. All scientific content, interpretation of results, and conclusions were developed and reviewed by the authors, who take full responsibility for the content of this publication.

### d. Declaration of Funding

This project was funded by the Marine Species Team, Department of Conservation, New Zealand.

### e. Conflict of Interest Disclosure

The authors have no known conflicts of interest.

### f. Permission to Reproduce Material From Other Sources

The authors confirm that all figures, tables, and other content included in this manuscript are original.

## AUTHOR CONTRIBUTIONS

Project conception KM; tagging KM, AJ, MB, CC, IJ, JC; data acquisition KM; data analysis GV, KM; methodology KM, AJ, MB, CC, IJ, JC; writing – draft KM, review all. All authors’ have read and agreed to publish this version of the manuscript.

## ACKNOWLEDGEMENTS

We acknowledge and thank iwi and hapu, who whakapapa (identify) to Rangaunu Harbour, for their support; Ngāi Takoto, Ngāti Kahu, and Te Whānau Moana/Te Rorohuri hapū of Ngāti Kahu. Local tamariki kura (school children) named two of the turtles Te Rito and Pumautahi and we commend them for their interest in honu conservation as young eco-warriors. Thanks also to all the amazing DOC Operations Teams around the moto (country), especially DOC Kaitaia staff, Auckland Zoo, and Sealife Kelly Tarlton’s staff for their long-term support for turtle conservation. We are also grateful for the GIS skills of Jodie Robertson and Peter Bradshaw (DOC), who updated the habitat classification map of Rangaunu Harbour from satellite and lidar images.

## Notes

### Competing Interest Statement

The authors have declared no competing interest.

