## Supplementary figures and images for "Green turtle satellite telemetry: identifying spatial movements, residency patterns, and habitat associations in Aotearoa New Zealand"

### Supplementary Figure 1

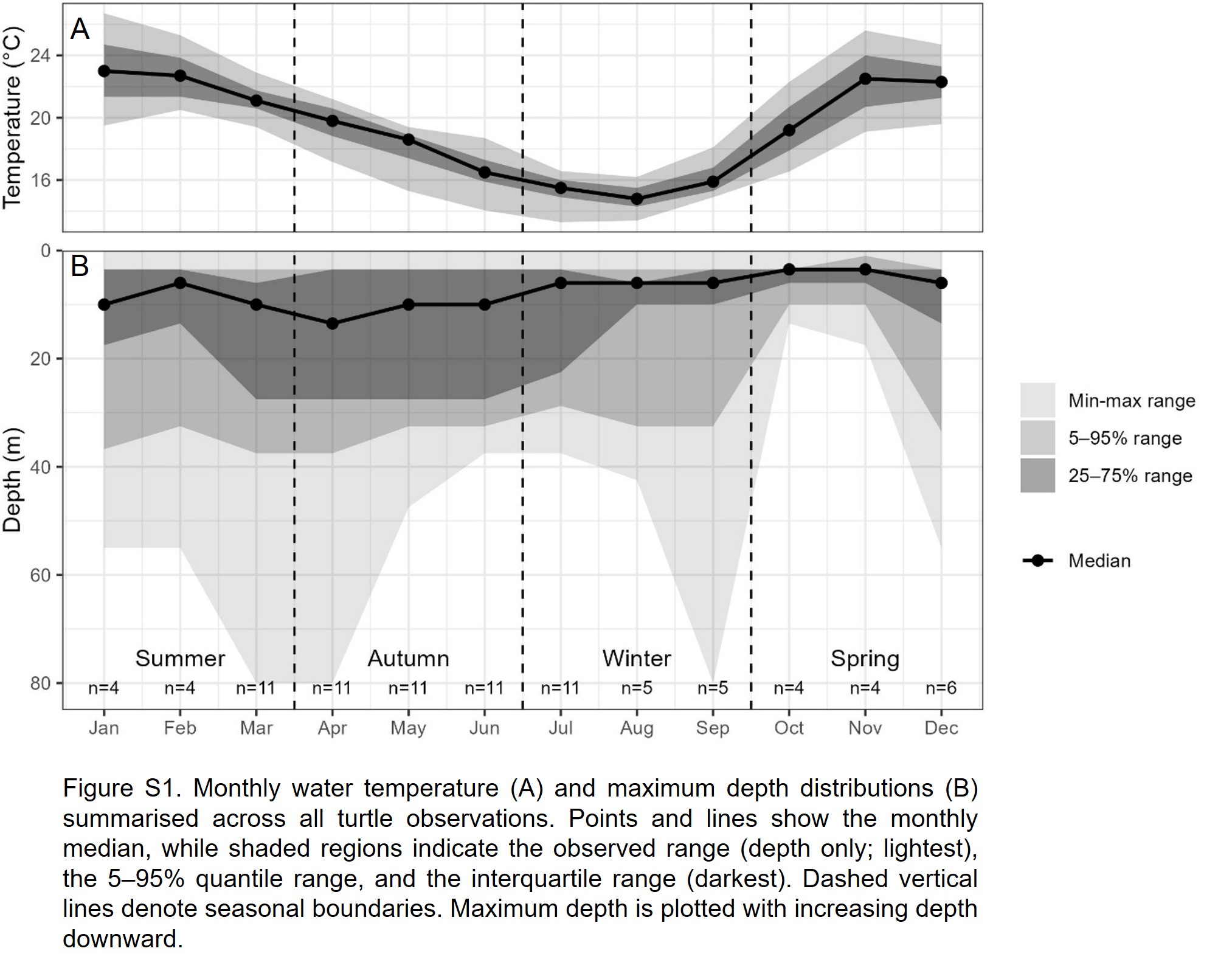
